# Inducible lipid storage and steatosis in the human choroid plexus associated with age and adiposity

**DOI:** 10.64898/2026.05.21.725559

**Authors:** N. Victoria Espericueta, Michael J. Neel, Yao Wang, Todd J. Soo, Parastou Porahang, Frances A. Goyokpin, Ryan S. Salehi, Sara Khan, Jasmine Lee, Nicolas B. Maramica, Guillermo Flores, Aditi Kulkarni, Sahil S. Plaha, Sam Ghahremani, William Huang, Quinton Smith, Peter Chang, Brett A. Johnson, Edwin S. Monuki

## Abstract

Cells that store lipids for other cells or organs can contain “giant” or large lipid droplets (LLDs) greater than 2 µm in diameter. In this study, human postmortem choroid plexus was evaluated for lipid droplets. Staining with hematoxylin and eosin (H&E), the lipophilic dye Oil red O, and anti-adipophilin antibodies established the presence of LLDs exceeding 10 µm in diameter in choroid plexus epithelial cells (CPECs). Manual annotation of H&E stains from 105 cases revealed a significant association between age and the percentage of CPECs containing LLDs (reaching up to 69%) and involving LLDs in our largest annotated category (>5 µm in diameter). The LLD association with age was replicated and extended to a total of 245 cases using a trained convolutional neural network, which further showed significant associations with body mass index at time of death (increasing with BMI), sex (higher in females >65 years old), and a near-significant association with Alzheimer’s Disease (lower in AD). Like HepG2 and derived hepatocytes, excess fatty acids in culture media readily induced LLDs and steatosis in human embryonic stem cell-derived CPECs. Akin to hepatocytes for the human body, we propose that CPECs store lipids for the human brain and become steatotic in the setting of excess adiposity.

## INTRODUCTION

Cells store fat in lipid droplet organelles, which consist of a triglyceride and cholesterol core surrounded by a phospholipid monolayer containing perilipins and other proteins (Gluchowski et al. 2017, Walther et al. 2017, Olzmann and Carvalho 2019, Kimmel et al. 2010). While most cells store lipid droplets smaller than one micron in diameter for use in their own energy production, membrane construction, and protection against lipotoxicity, specialized cells such as enterocytes, adipocytes, mammary epithelial cells, hepatocytes and hepatic stellate cells contain “giant” lipid droplets greater than 2 microns across (Walther et al. 2017). The latter cells store lipids for other cells and organs – enterocytes after fat-heavy meals for postprandial storage (Beilstein et al. 2016, Xiao et al. 2019, Robertson et al. 2003), adipocytes as a long-term energy store (Suzuki et al. 2011, Richard et al. 2020), mammary epithelial cells for milk fat globule production to nourish the young (Russell et al. 2007, Cohen et al. 2015), hepatocytes when adipocyte storage capacity is saturated (Heeren and Scheja 2021), and hepatic stellate cells as long term retinoid (Vitamin A) storage (Blaner et al. 2009). Excess lipid accumulation in hepatocytes, which normally esterify fatty acids and package lipoproteins for bloodborne whole-body distribution, can become a pathological situation proceeding through microvesicular and macrovesicular steatosis to non-alcoholic fatty liver disease (NAFLD), non-alcoholic steatohepatitis (NASH), and liver cirrhosis (Marchisello et al. 2019, Heeren and Scheja 2021, Scorletti and Carr 2022).

Much as the liver conditions the blood, the choroid plexus (ChP) produces and conditions the cerebrospinal fluid (CSF) by pumping water, synthesizing and secreting proteins, and transporting various nutrients into and toxins out of the ventricles (for a recent review, see Saunders et al. 2023). While creating the blood-CSF barrier, the ChP also provides a gateway for immune cell entry into the brain (Wewer et al. 2011; Xu et al. 2024), and because it has free access to the circulatory system through fenestrated capillaries, it is a potential therapeutic target for peripherally administered drugs (Dabbagh et al. 2022, Thompson et al. 2022).

Human ChP epithelial cells (CPECs) contain vesicles that appear as empty vacuoles after staining with hematoxylin and eosin (H&E) and that increase in abundance with age (Dunn and Kernohan 1955). At least a fraction of CPEC vesicles stain with lipophilic dyes, suggesting the presence of lipids (Dunn and Kernohan 1955), and apparent lipid bodies were described in early electron microscopic studies of human ChP (Oksche et al. 1969, 1972). CPECs in frogs (Paul 1970) and rabbits (Meek 1907, Weindl et al. 1969, Marinetti et al. 1971) have lipid vesicles sufficiently large to qualify as giant lipid droplets, as do CPECs of mice fed a high-fat diet (Denes et al. 2012), but the existence of giant lipid droplets in human CPECs remains obscure.

Our laboratory is engaged in an extensive and systematic survey of various age-related changes to the ChP and their potential associations with disease and other clinical conditions, drawing from repositories of postmortem tissue at the University of California, Irvine and combining both manual and machine learning approaches. In this study, we defined the CPEC vesicles/vacuoles to be giant or large lipid droplets (LLDs), and found that LLDs increase in abundance (% CPECs affected) with both age and body mass index (BMI). To mimic high BMI, excess fatty acids were then applied to human embryonic stem cell-derived CPECs, which readily formed LLDs and became steatotic.

## RESULTS

### Large vacuoles in H&E sections are lipid droplets

We have noted that vacuoles in CPECs of H&E-stained ChP vary considerably from case to case in our tissue repository, presenting similarly to hepatosteatosis (vacuolar changes in the hepatocytes known as micro- and macrovesicular steatosis) in some instances. For example, CPECs from the very high BMI individual illustrated in Fig. 1A have both large and small vacuoles, often including multiple vacuoles per cell.

**Fig. 1.**
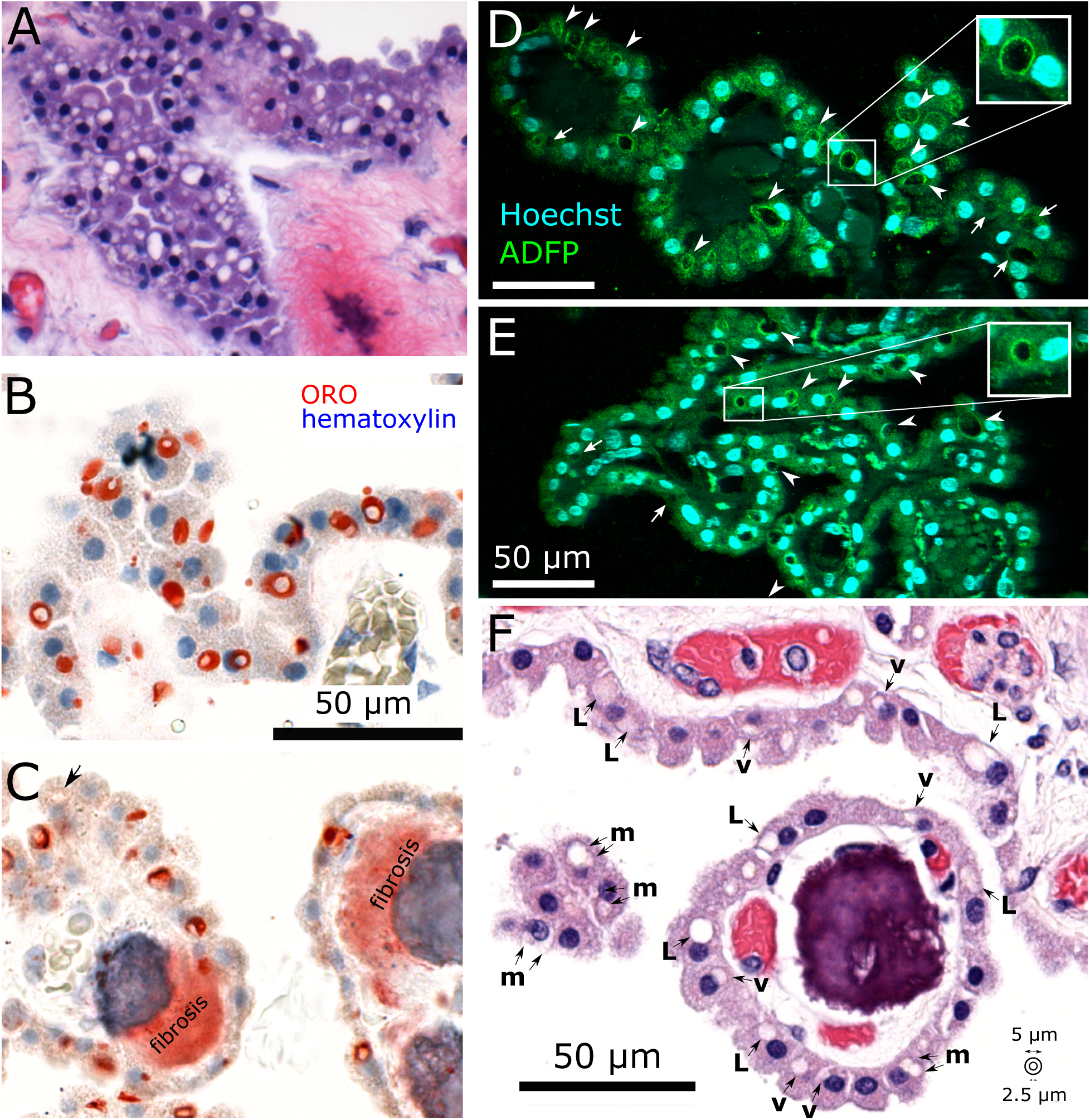
Lipid droplets are present in postmortem human ChP. **A.** Image from an H&E-stained section of ChP from an individual with high BMI (185324) showing extensive vacuolation in CPECs. **B,C.** Oil red O (ORO) staining of lipid droplets in human CPECs. Twelve-micron cryosections of formalin-fixed choroid plexus were stained with ORO and counter-stained with hematoxylin. Images of two cases (**B:** 185324; **C:** 586728) were chosen to illustrate the range of sizes and numbers of droplets in different CPECs and cases. **C**. Areas of fibrosis also bound the ORO; the adjacent blue hematoxylin lake indicates mineralization of part of the fibrotic area. Similar results were obtained for three other cases. **D,E.** Immunofluorescent staining of membranes of CPEC lipid droplets with an antibody to adipophilin (ADFP) on 5-µm FFPE sections. Well-labeled droplets are indicated with arrowheads, while poorly labeled or unlabeled vacuoles are indicated with arrows. Images of two cases with abundant vacuoles in H&E-stained sections (**D:** 185324; **E:** 822832) are shown to illustrate the generality of the finding. The image in **D** is from the same case as Fig. 1A. Similar results were obtained for ChP from six other cases. Images are maximal z-projections from confocal microscopy. **F.** Lipid droplets annotated as vacuoles in 5-µm FFPE sections stained with H&E. For quantification of the effects of age and body mass index, CPECs were annotated as containing either large vacuoles (L, > 5 µm across), multiple vacuoles (m) > 2.5 µm across, or vacuoles (v) between 2.5 and 5 µm across.

Suspecting that such vacuoles may represent lipid droplets, fixed postmortem specimens of atrial ChP were cryosectioned and stained with the lipophilic dye Oil red O (ORO). Indeed, lipid droplets within the CPECs were readily visualized (Fig. 1B,C). For five cases, we quantified the percent of CPECs affected by lipid droplets using Oil red O-stained sections and compared them to results obtained on separate H&E sections. There was a high correlation between the two staining methods for these five cases (r = 0.95; Fig. S4).

The droplets varied in abundance and size both across a single ChP and across ChP from different individuals, often exceeding 5 µm in diameter. In a single histological section, some cells appeared to lack droplets, while others appeared to contain more than one. Vacuoles without ORO staining also were observed in CPECs (arrow in Fig. 1C), but these were present at only 0.019 times the frequency of CPECs with clear ORO staining (average of 2 annotators, range of 0.015-0.023). Many lipid droplets appeared to have a hollow core, although it is unclear whether this feature is natural or artefactual (e.g., lipid loss from LLD cores due to the polyethylene glycol ORO solvent and/or fixation prior to cryosectioning). ORO also stained stromal fibrosis to some extent, although not as intensely as LLDs (Fig. 1C).

Lipid droplet organelles are characterized by perilipin membrane proteins, and the most common perilipin across cell types appears to be adipophilin (aka ADFP, ADRP, perilipin 2, PLIN2; Itabe et al. 2017). Immunofluorescent staining of FFPE sections of atrial ChP with an antibody to adipophilin resulted in labeling of the perimeters of vacuoles/vesicles within CPECs (Fig. 1D & E, arrowheads). There also were indications of vacuoles without associated immunoreactivity (Fig. 1D & E, arrows), which could reflect the dynamic nature of adipophilin’s association with lipid droplets (Straub et al. 2008, Kimmel et al. 2010), given the low frequency of vacuoles lacking ORO staining, as mentioned above.

### Large lipid vacuoles in CPECs are associated with aging

We selected four cases, measured the areas of vacuoles exceeding 2.5 µm along their longest axis, and calculated the diameters of circles giving these areas. We found that vacuoles could exceed 10 µm in diameter and that the four cases differed significantly in median diameter (Kruskal-Wallis statistic = 211.5, p < 0.0001) (Fig. 2). For example, almost half of the vacuoles in case 783896 exceeded 5 µm in diameter, whereas vacuoles of this size were rarer in case 763917. On the other hand, the distribution of sizes in cases 447764 and 164179 were statistically indistinguishable from each other, but different from both case 783896 and case 763917 [see Table 1 for pairwise nonparametric comparisons of median size (Mann-Whitney U) and size distribution (Kolmogorov-Smirnov). Similar results were obtained by three independent annotators, who differed most in their assessment of smaller vacuoles (Fig. S5, Table S3), likely due to lesser contrast with surrounding cytoplasm as their diameters would be smaller than the section thickness.

**Fig. 2.**
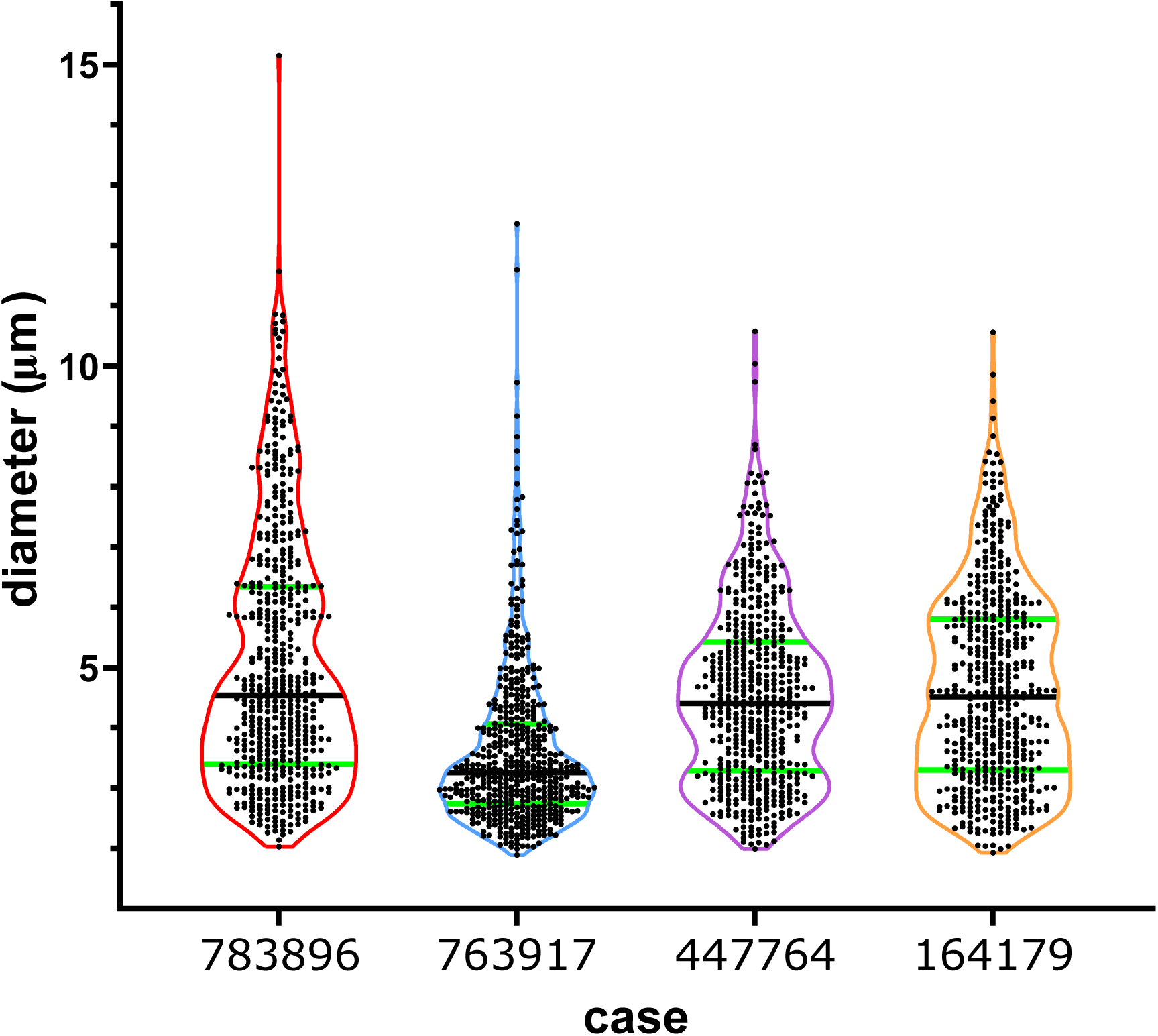
Lipid droplets differ in size across different cases and can exceed 10 microns in diameter. Violin plots displaying the sizes of vacuoles traced in areas randomly selected from H&E-stained whole slide images of ChP from four cases. The areas of vacuoles exceeding 2.5 µm along their major axes were measured, and the diameters of the circles that would result in those areas were calculated. Entire tiles were sampled until at least 500 vacuoles had been analyzed (524 for 783896, 512 for 763917, 518 for 447764, and 510 for 164179). Each point represents a vacuole, median sizes are indicated with a black line and quartiles are shown in green. Table 1 shows results of statistical analyses indicating that cases differed significantly in size distributions. Results shown here were obtained by a single annotator (BAJ); similar results were obtained by two additional annotators (Table S2).

**Table 1.** Statistically significant differences (*) in sizes (Mann-Whitney U tests) and size distributions (Kolmogorov-Smirnov tests) between pairs of cases shown in Fig. 2 (783896, n=524; 763917, n=512; 447764, n=518; 164179, n=510). Values shown here represent a single manual annotator (BAJ).

| **Mann-Whitney U** |  |  |  |  |  |  |
| --- | --- | --- | --- | --- | --- | --- |
|  | 763917 |  | 447764 |  | 164179 |  |
|  | U | p | U | p | U | p |
| 783896 | 72506 | <0.0001* | 120926 | 0.0023* | 121588 | 0.012* |
| 763917 |  |  | 79688 | <0.0001* | 78571 | <0.0001* |
| 447764 |  |  |  |  | 128328 | 0.43 |
| **Kolmogorov-Smirnov** |  |  |  |  |  |  |
|  | 763917 |  | 447764 |  | 164179 |  |
|  | D | p | D | p | D | p |
| 783896 | 0.354 | <0.0001* | 0.152 | <0.0001* | 0.110 | 0.0038* |
| 763917 |  |  | 0.350 | <0.0001* | 0.338 | <0.0001* |
| 447764 |  |  |  |  | 0.081 | 0.071 |

Our repository of FFPE ChP from the atrium of the lateral ventricle enabled quantitative analysis of vacuoles in H&E-stained sections. We selected 105 cases to analyze manually. Given the differences in sizes and numbers of lipid droplets across different cases and CPECs, we annotated CPECs as belonging to one of four vacuole categories (unaffected, single vacuole between 2.5 and 5 µm across, single vacuole exceeding 5 µm, or multiple vacuoles > 2.5 µm) (Fig. 1F). Data for these cases are tabulated in Table S1. Statistical analysis on the three vacuole categories showed a strong effect of age, but not of source or sex (Table 2). Univariate analyses isolated the age effect to only the largest vacuole category (Fig. 3A, Table 2).

**Fig. 3.**
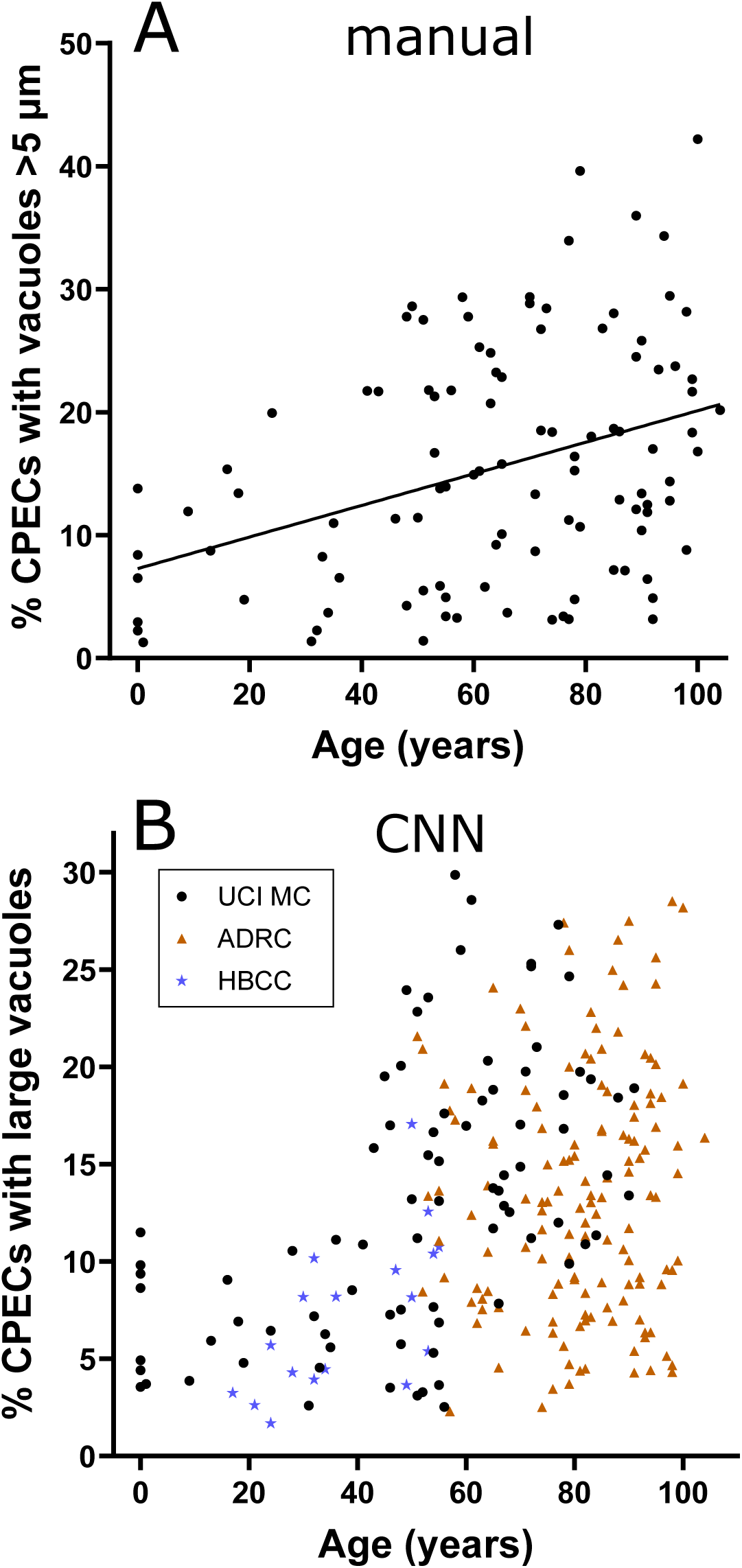
Large lipid droplets increase in association with age. The percentage of CPECs containing vacuoles >5 microns across increased significantly both in the manually annotated cases (**A**, n=105) and in the cases annotated by convolutional neural networks (**B**, n=245). The trendline in **A** is the result of a simple linear regression. In **B**, different plot symbols are used to elucidate the effect of different tissue sources. **ADRC**, Alzheimer’s Disease Research Center; **UCIMC**, autopsy service at the UC Irvine Medical Center; **HBCC**, Human Brain Collection Core of the NIMH.

**Table 2.** Statistical analysis of effects of age, source, and sex on prevalence of vacuole categories (manual annotations). Boldface denotes p < 0.05.

| p-values |  |  |  |  | n |  |
| --- | --- | --- | --- | --- | --- | --- |
|  | Multivariate | Univariate |  |  |  |  |
|  |  | single, > 5 microns | single, 2.5 – 5 microns | multiple, >2.5 microns |  |  |
| Age | <0.001 | <0.001 | 0.742 | 0.410 | Total | 105 |
| Sex | 0.784 | 0.585 | 0.439 | 0.949 | Male | 55 |
| Source | 0.403 | 0.101 | 0.544 | 0.221 | Female | 50 |

### Deep machine learning reveals an association between large lipid vacuoles and adiposity (BMI)

Multiple human annotators contributed hundreds of hours to collect the data presented above. To enable efficient extensions of this work and to eliminate effects of interobserver variance, we developed a two-stage, deep learning, convolutional neural network (CNN) approach (detection of CPECs followed by binary classification for large vacuole presence; Fig. S3). The coordinates of CPECs with and without vacuoles in the largest category (>5 µm diameter) from the manual analysis were used as training data, given that this category was more sensitive to age effects and showed lesser variance between manual annotators.

After training, the CPEC detection model achieved a median intersection over union (IOU) of 0.71 for boxes drawn around CPECs compared to manual annotations. When compared to 3 whole slide images fully annotated for CPECs, the model had a mean CPEC detection sensitivity, precision, and false discovery rate of 0.84, 0.87, and 0.13, respectively (Table S4). After training of the large vacuole classifier, the model had an accuracy, sensitivity, and precision of 0.89, 0.62, and 0.60, respectively, and an area under the receiver operating characteristic curve (AUROC) of 0.87 (Fig. S6). The results of the models, which assessed vacuoles across entire whole slide images, correlated well with the manual data, which assessed only fractions of each section (Fig. S7).

Using the deep learning model, we expanded our analysis to a total of 245 cases from three different sources (UCIMC, UCI ADRC, and HBCC). The HBCC samples included material from temporal and frontal horns of the lateral ventricle whereas samples from the other sources were restricted to the atrium; however, we detected no systematic differences across these three locations in a repeated measures ANOVA (n=13 cases with specimens from all three locations; Fig. S8).

The network approach on the extended dataset recapitulated the significant effect of age. Tissue source also had a significant effect (Fig. 3B, Table 3), which is not surprising given the systematic differences in age across the three sources. These sources also varied in postmortem intervals and fixation procedures (see Methods), but postmortem interval (PMI) was not found to be significant for the reduced number of cases for which PMI data were available (n=172, p=0.52)

**Table 3.**
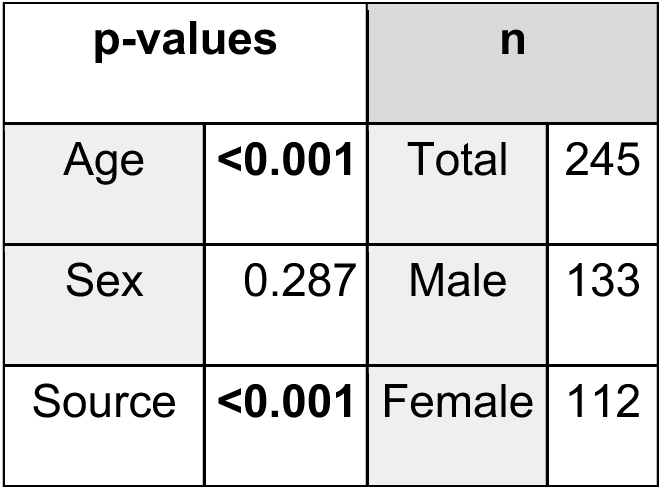
Statistical analysis of large (>5 micron) vacuole prevalence (convolutional neural network annotations). Boldface denotes p < 0.05.

| p-values |  | n |  |
| --- | --- | --- | --- |
| Age | **<0.001** | Total | 245 |
| Sex | 0.287 | Male | 133 |
| Source | **<0.001** | Female | 112 |

Many of our cases were from the UCI ADRC, and therefore sporadic Alzheimer’s Disease (AD) was well represented in the population. Elderly cases from UCIMC also provided both controls and AD cases. Overall, we extracted 151 total cases ≥ 65 years old (74 AD, 77 non-AD; 86 male, 65 female; 29 UCIMC, 122 ADRC). Sex (p=0.031, females having more large vacuoles) and source (p=0.048, ADRC having fewer large vacuoles) were identified as significant factors in this older population, while AD approached significance (p=0.061, AD cases having fewer large vacuoles). Age had no effect in individuals above 65 years (p=0.683).

The resemblance of steatotic CPECs to hepatocytes raised the possibility of an association with excess body fat. To explore this possibility, we analyzed the large lipid vacuoles with respect to body mass index (BMI) at autopsy, drawing from the UCIMC and HBCC cases that included BMI data (84 total cases; 51 male, 33 female; 66 UCIMC, 18 HBCC). BMI data (height and weight) were unavailable for ADRC cases, and UCIMC cases < 1 year old were excluded. Higher BMI was associated with a higher prevalence of large CPEC vacuoles (p=0.01, Fig. 4A). Age (p < 0.001) was also a significant factor in this population (Fig. 4B), as was source (p=0.007), while sex was without an impact in this generally younger group (p=0.662). When the highest BMI (57.6) index case illustrated in Fig. 1A was excluded, a p value of 0.046 was obtained, consistent with a considerable association between BMI and large lipid vacuoles in the remaining data set.

**Fig. 4.**
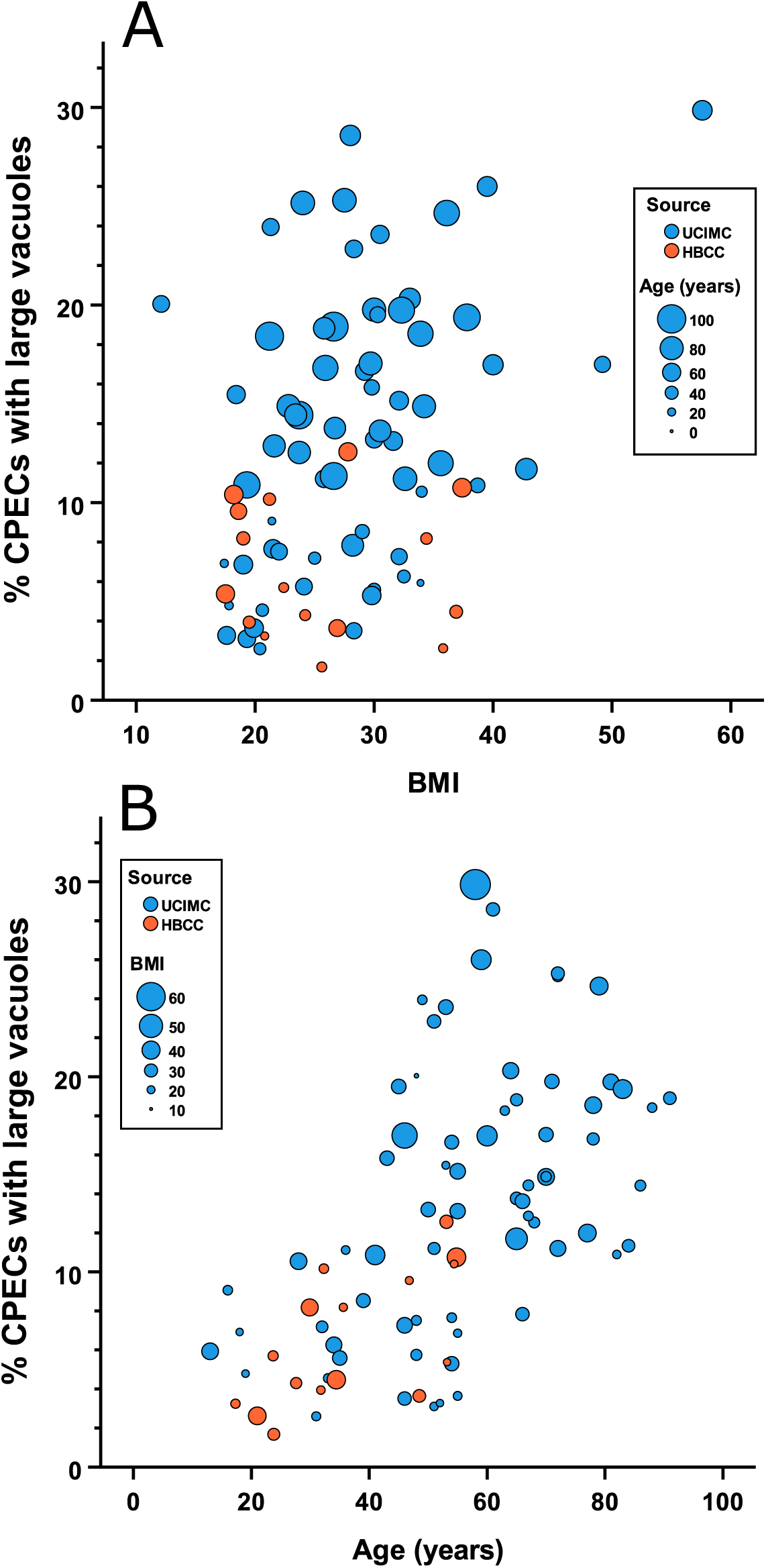
Large lipid droplets increase in association with BMI as determined using convolutional neural network analysis. Data are plotted in two ways to illustrate the combined effects of BMI, age, and tissue source. **A**. BMI is plotted along the x-axis while age is represented by plot symbol size. **B**. BMI is represented by plot symbol size, while age is plotted along the x-axis. **UCIMC**, autopsy service at the UC Irvine Medical Center; **HBCC**, Human Brain Collection Core of the NIMH.

### Excess fatty acids induce steatosis and large lipid droplets in stem cell-derived CPECs

To investigate directly the associations between excess fat, large lipid droplets, and steatosis, we established an *in vitro* model using human ESC-derived CPECs (dCPECs) which was based on studies using hepatocytes (Cui et al. 2010, Rida et al. 2023, and Wang et al. 2026). After confirming with iPSC-derived hepatocyte cultures (Fig. S9), we treated 88-day dCPECs with 600 µM oleate coupled to albumin (BSA), 600 µM palmitate coupled to BSA, or BSA alone for 48 hours. Using LipidTOX for visualization, steatosis was qualitatively apparent in oleate and palmitate-treated dCPECs compared to BSA controls, while LLDs were most obvious in the oleate-treated cultures (Fig. 5A). Quantitative analysis demonstrated that both oleate and palmitate increased lipid load per cell compared to BSA controls (Fig 5B; Kruskal-Wallis statistic = 41.04, p < 0.0001). Size distribution varied across conditions (Kruskal-Wallis statistic = 303.8, p < 0.0001). Lipid droplets >2 µm in diameter were present in all three conditions, but only oleate-treated dCPECs had lipid droplets >5 µm in diameter and skewed larger overall compared to vehicle or palmitate-treated dCPECs (Fig. 5C).

**Fig. 5.**
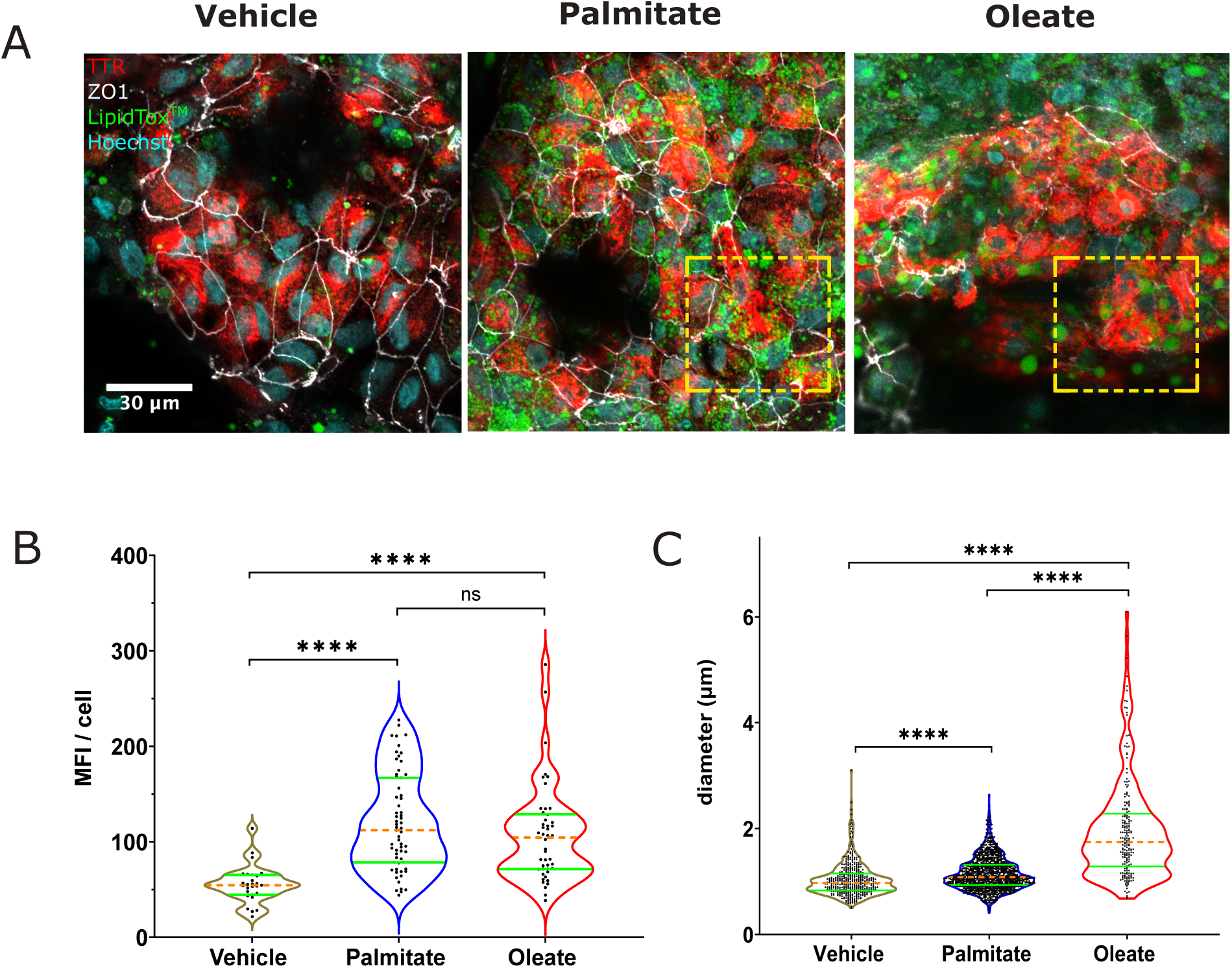
iPSC-derived CPECs can be induced to form large lipid droplets. **A.** Five-micron 40X confocal images of CPECs treated with vehicle (BSA), palmitate-BSA, or oleate-BSA were stained for TTR (red), ZO1 (white), LipidTOX^TM^ (green), and Hoechst (blue). Yellow dashed boxes highlight the size difference in palmitate-induced lipid droplets versus oleate-induced lipid droplets. Scale bar: 30µm. **B.** Violin plots displaying mean fluorescence intensity (MFI) of LipidTOX^TM^ per cell across treatment conditions. **C.** Violin plots displaying lipid droplet diameter sizes across treatment conditions. Mann-Whitney tests were performed for pairwise comparisons; ****, p<0.0001; ns, not significant.

## DISCUSSION

Our results have implications for both the physiology and the pathology of human CPECs. Oil red O staining and ADFP immunostaining revealed the presence of large (>10 µm in diameter) lipid droplets in human CPECs, indicating a similarity with cell types storing lipid for delivery to other cells. For CPECs, the likely recipients are other brain cells, which use fatty acids for membrane integrity and fluidity, and to protect against inflammation (Qadir et al. 2026). Indeed, lipid-carrying apolipoproteins, especially clusterin (CLU or APOJ) and apolipoprotein E (APOE), are among the genes most highly transcribed by CPECs (Marques et al. 2011, Janssen et al. 2013, Lun et al. 2015, Bergen et al. 2015), suggesting that an important function of CPECs is to package lipoproteins for secretion into the CSF (Fujiyoshi et al. 2007, Achariyar et al. 2016, Kaiser and Bryja 2020), much as hepatocytes deliver lipoproteins to the blood (Feingold 2021).

We have demonstrated that stem cell-derived CPECs take up albumin-associated fatty acids from the surrounding media and package them into lipid droplets *in vitro*. Circulating triglyceride-rich lipoproteins and free fatty acids (FFA) are the sources of fatty acids for peripheral tissues. Lipoprotein lipase (LPL) hydrolyzes triglycerides from lipoproteins at the capillary lumen, releasing free fatty acids that subsequently bind albumin prior to cellular uptake. A recent study (Song et al. 2025) identified CPECs as a source of LPL in choroid plexus capillaries, where it is transported across the endothelium and anchored at the luminal surface by GPIHBP1. The authors proposed that the resulting fatty acids are subsequently taken up by CPECs. Our findings extend this model by demonstrating that CPECs internalize the albumin-bound fatty acids and store them in lipid droplets that can reach notable diameters.

Consistent with findings in derived CPECs, we have found that the number and size of human CPEC vacuoles in H&E-stained sections are correlated with BMI, which may at least partially explain previously reported associations between obesity or waist circumference and ChP volume and microstructure as measured using MRI (Alisch et al. 2022). An association between BMI and human CPEC lipid droplets is also consistent with the qualitative detection of lipid droplets in lateral ventricle CPECs of mice fed a high-fat Paigen of Western diet (Denes et al. 2012).

In the liver, large lipid droplet accumulation is pathological, and like lipid droplets in the human ChP, hepatosteatosis and NAFLD are correlated with both age (Hilden et al. 1977, Younossi et al. 2016) and BMI (Loomis et al. 2016, Hagström et al. 2016, Wang et al. 2016, Fan et al. 2018, Pasanta et al. 2018, Pang et al. 2019, Lu et al. 2023, Tokutsu et al. 2023). The accumulation of large lipid droplets in *in vitro* models of hepatosteatosis is correlated with changes in transcription and metabolism (Gunn et al. 2020), including compromised electron transport chain function (Sinton et al. 2020), as well as increased cell stiffness, nuclear deformation, cytoskeletal network disruption, and impaired mechanosensing (Chin et al. 2020, Loneker et al. 2023). In addition, livers showing steatosis perform poorly after transplantation (Chu et al. 2015). Less research has been conducted on the possible relationships between obesity and ChP functions, although one study found an association between high BMI and choroid plexus barrier dysfunction (Brettschneider et al. 2005). Obese Zucker rats have been reported to have higher aquaporin 1 water channel expression in the ChP (Udall et al. 2017), and obese mice fed a Western diet display changes in transcription of genes encoding junctional and transport proteins in endothelial cells of ChP fenestrated capillaries, as well as reduced ChP uptake of dyes from blood (Bondareva et al. 2022). Along with various other aging-related and likely detrimental changes to the choroid plexus (Dunn and Kernohan 1955), CPEC steatosis may contribute to age-related reduction in CSF production, putting the brain at risk for degenerative diseases and cognitive decline (Attier-Zmudka et al. 2019).

In our analysis, vacuolation of CPECs was lower in AD, approaching statistical significance, which contrasts with a previous report of increased ChP lipid associated with AD severity and AD risk genes (Yin et al. 2019). However, much of the Oil red O staining analyzed in the Yin et al. study appeared to have been present in stroma, possibly corresponding to the staining of fibrotic areas in stroma that we also have observed, but were not analyzed in our study (Fig. 1C). It is well established in cohorts across the globe that losses of body weight and BMI accompany the later stages of AD, attributable to a reverse causal association wherein AD patients have reduced caloric intake (Albanese et al. 2013, Singh-Manoux et al. 2018, Kivimaki et al. 2018, Ciciliati et al. 2021). Thus, the near-significant reduction of lipid vacuoles at death in our AD cases may be related to a lower BMI in this group, for which BMI data at death was not available. In a study collecting data at autopsy, as we have done, BMI also was lower in older males than in older females (Ciciliati et al. 2021), which may explain why males had fewer large lipid vacuoles in our older cohort.

Our results involving postmortem material from hospital autopsies and neurological disease specimens may be confounded in part by short-term factors that can affect vacuolation, which may cloud even stronger relationships with age and BMI. For example, rodents exposed to tertiary amines and/or cationic amphiphilic compounds quickly develop CPEC vacuoles (Benitz and Kramer 1968, Levine and Sowinski 1977, Frisch and Lüllmann-Rauch 1979), which may be related to lipidosis (Frisch and Lüllmann-Rauch 1979); hypobaric hypoxia can induce CPEC vacuolation (Wang and Kaur 2001); and increased CPEC vacuolation has been reported following head trauma (Rand and Courville 1931). In cell culture models of hepatocytes, adipocytes, enterocytes, and mammary epithelial cells, lipid droplets can be induced and reversed within hours or days following manipulation of culture media to alter lipid uptake or biogenesis by adding or removing fats, sugars, and other energy sources (Gunn et al. 2020, Wolins et al. 2005, Lagrutta et al. 2017, Lyall et al. 2018, Cohen et al. 2015, Sinton et al. 2020, Bouchoux et al. 2011). However, short-term effects of reduced blood lipids or sugars on CPECs due to illness may be partially offset by continued availability of lipids from adipocyte stores that are reflected in BMI at autopsy.

Using the manual annotations of H&E vacuoles as training data, we developed CNNs that effectively replicated, refined, and expanded relationships between lipid droplets and both age and BMI. Deep learning machine models to analyze postmortem tissue for ChP features such as lipid droplets and Biondi body inclusions (Neel et al., in preparation) will provide a high-throughput process by which we can identify other age and disease-related changes that may affect brain homeostasis.

## ACKNOWLEDGEMENTS

We thank Dr. Joni Ricks-Oddie and Dr. Farideh Dehkordi-Vakil for their expert statistical advice, and Jae Shin for early contributions to the vacuole annotation procedures. We also thank UCI neuropathologists and technicians at the Experimental Tissue Resource and UCI ADRC Neuropathology Core for their assistance in procuring and processing tissue, staining, and slide scanning. Some of the tissue used in this research was obtained from the Human Brain Collection Core, Intramural Research Program, NIMH (http://www.nimh.nih.gov/hbcc) with the assistance of Dr. Stefano Marenco and Dr. Pavan Auluck. This work was supported by the following NIH awards: P50AG016573 and P30AG066519 (UCI ADRC), T32AG073088 (N.V.E. and M.J.N.), T32AG000096 (M.J.N.), and R21MH109036 and R21AG064640 (E.S.M.). This work was also supported by the following awards from the California Institute for Regenerative Medicine (CIRM): EDUC4-12822 (Y.W.), EDUC2-12638 (J.L.), EDUC5-13647 (G.F.), and RN2-00915-1 (E.S.M.). Finally, we thank the tissue donors and their families for making this research possible.

## METHODS

### Postmortem tissue

Specimens of ChP were obtained from the UC Irvine Medical Center (UCIMC) autopsy service, the UCI Alzheimer’s Disease Research Center (UCI ADRC), and the Human Brain Collection Core (HBCC) at the NIMH (Supplemental Table 1). At UCI, the ChP was taken from known locations of the lateral ventricle (mostly the glomus of the ChP, or glomus choroideum, in the atrium or trigone of the lateral ventricle) during cutting of fixed brains. In addition to systematic differences in median postmortem intervals (ADRC 5.3 hr < HBCC 31.5 hr < MC 124.5 hr), the autopsy service fixed brains in formalin, whereas the ADRC used freshly prepared 4% paraformaldehyde. At the HBCC, lateral ventricle ChP from the atrium, frontal horn, or temporal horn of the lateral ventricle was dissected from frozen brain slices and shipped on dry ice to UCI, where they were drop-fixed at room temperature in 10% formalin for 24 hours with gentle agitation (Electron Microscopy Sciences, 15740-01). For all cases, tissue specimens were embedded in paraffin, sectioned at 5-µm thickness, and then either stained using hematoxylin and eosin (H&E) through a UCI core facility (Experimental Tissue Resource) or supplied unstained for immunostaining procedures. A single UCIMC case, with a diagnosis of cystinosis, was excluded from the study due to unusual dark inclusions within CPEC vacuoles that complicated discrimination of lipid droplets. For some cases, fixed samples of choroid plexus were supplied intact by UCI neuropathologists and stored in phosphate-buffered saline containing 0.02% sodium azide before being dissected, mounted in O.C.T. compound, and sectioned at 12-µm thickness in a cryostat for Oil red O (ORO) staining.

### Stem cell culture and cell culture derivations

H1 embryonic cell line (ESCs; WiCell Research Institute, Madison, WI) or KOLF2.1J cell line (iPSCs; The Jackson Laboratory, Bar Harbor, ME) were cultured and differentiated into CPECs or hepatocytes respectively, as previously described (Masters et al. 2025; Wang et al. 2026), with the following modifications: 1) After reaching confluence, ESCs were passaged using E8 media with CEPT cocktail (Chen et al. 2021) rather than ROCK inhibitor alone; and 2) during CPEC differentiation from ESCs, BMP4 exposure was reduced from 15 to 10 days to reduce cell death.

### Histology and immunocytochemistry

ORO staining was performed by the UCI Experimental Tissue Resource using polyethylene glycol solvent and hematoxylin counterstain. Immunostaining employed deparaffinized sections (xylenes followed by a descending series of ethanol concentrations), antigen retrieval in 10 mM sodium citrate, pH6 in a vegetable steamer for 20 minutes, blocking in 5% donkey serum and 0.3% Triton X-100, and overnight incubation with rabbit anti-adipophilin polyclonal antibody (1:200 dilution of Invitrogen #PA1-16972 in 1% donkey serum and 0.3% Triton X-100) at 4°C. Antibodies were detected using 1:200 Alexa 488-donkey anti-rabbit secondary antibody (Invitrogen #A32790) for 1 hour at room temperature followed by counterstaining with Hoechst 33342 (Invitrogen). Images were acquired using an Olympus FV3000 laser-scanning confocal spectral inverted microscope and a Plan-Apo 40X /1.25 oil objective.

Derived CPEC cultures were gently washed twice with Dulbecco’s Phosphate Buffered Saline (DPBS), fixed with 4% paraformaldehyde at room temperature for 15 minutes, then washed three times with DPBS. Immunocytochemistry consisted of a 1-hour blocking incubation in 5% donkey serum and 0.3% Triton X-100, and overnight incubation with sheep anti-transthyretin and rabbit anti-zonula occludens-1 polyclonal antibodies (1:3000 dilution of Abcam #AB9015 and 1:1000 dilution of Invitrogen #61-7300 in 1% donkey serum and 0.3% Triton X-100) at 4°C. Derived hepatocyte cultures were processed the same but using rabbit-ASGR1 (1:500 dilution of Invitrogen #PA5-52885 1% donkey serum and 0.3% Triton X-100) instead. Primary antibodies were detected using 1:500 Alexa 555-donkey anti-sheep, Alexa 647 or 555-donkey anti-rabbit secondary antibodies (Invitrogen #A-21436, #A-31573, and #A-31572) for 1 hour at room temperature followed by counterstaining with Hoechst 33342 (Invitrogen #H3570) and 30 minute incubation with HCS LipidTOX™ (1:200 of Invitrogen #H34475). Images were collected using an Olympus FV3000 laser-scanning confocal spectral inverted microscope. Image acquisition was performed blinded to the lipid droplet fluorescence.

### Manual annotations

#### H&E vacuole annotations

H&E-stained sections were scanned into whole slide images at 40X using an Aperio Versa 2 scanner in the ADRC Neuropathology Core. Image files were converted to svs format using Aperio ImageScope v12.4 software, subdivided into 280.57-µm x 280.57-µm tiles, and down-sampled by a factor of 2 with a final resolution of 0.274 µm/pixel. Image tiles were further converted, in random order, to HDF5 stacked formats compatible with a custom-designed software client. This client presented the tiles to annotators in two image windows representing different magnifications. The lower magnification window was used to locate CPECs that contained vacuoles and to use a mouse and keystrokes to annotate CPECs depending on the category of vacuole they contained. The higher magnification window superimposed a reticle involving two concentric circles, corresponding to 2.5 and 5 µm diameters, and was used to classify the vacuoles as either meeting the minimal size criterion only (2.5-5 µm) or belonging to the larger category (>5 µm). Another category was used to indicate that CPECs contained more than one vacuole exceeding the minimal dimensions. CPECs lacking vacuoles below the minimum were separately annotated. In preparation for training a neural network to identify CPECs, a visible nucleus was required for all CPECs except for those containing the larger category of vacuoles, which often displaced and distorted the nuclei. Annotators completed evaluating all CPECs in a tile and then exported the results once completion was signaled based on stability of coefficients of error, as indicated in the legend to Fig. S1. These criteria led to the evaluation of 1955 +/- 739 (mean +/- standard deviation, range 726-4328) CPECs per case. In addition to quantifying vacuole and CPEC numbers, the software client recorded coordinates of the CPEC nuclei in each category. These coordinates were used to train convolutional neural networks (CNNs) first to recognize CPECs and then to classify them with respect to possessing vacuoles larger than 5 microns (see below under Deep learning models). Vacuoles were defined as objects with smooth (slightly elliptical or circular) contours that appeared lighter than the surrounding CPEC cytoplasm, but not necessarily transparent. Crescent-shaped spaces adjacent to nuclei (Golgi apparatus), almond-shaped gaps between CPECs (paracellular space), and vacuoles enclosing nuclear or other hematoxylin-stained material were explicitly omitted from the annotations. Annotators included seven undergraduate students as well as a senior project scientist (BAJ) (see Table S1). Initial training on CPEC and vacuole classification was given by a board-certified neuropathologist (ESM) followed by live interactive demonstrations, video recordings, illustrated manuals, and live regular feedback sessions with the project scientist. Annotations were conducted blind; annotators were unaware of any case information, variable, or covariable including age, sex, AD status, other diagnoses, or postmortem interval.

After training, all annotators completed analysis of the same set of tiles from three cases. The individual results, means, and standard deviations are shown in Fig. S2. The range of results across annotators was considerable; nevertheless, cases could be distinguished statistically despite the variance (Table S2), increasing our confidence in proceeding with the data analysis across cases analyzed by the different researchers. Replication of the results with a single machine-learning model further solidified confidence.

#### Lipid uptake annotations

Confocal Z-stacks of derived CPECs were acquired at 1 µm intervals at 40X objective using the Olympus FV3000 confocal microscope, and ImageJ was used to generate TIFF files of maximum intensity projections from five optical slices (5 µm total depth). TIFF files were imported onto QuPath (v.0.5.0; Bankhead et al. 2017) for tracing and analysis. For blinding purposes, individual CPECs were defined based on nuclear signal, membranous ZO1 and cytoplasmic TTR (blue, red, and far-red channels, respectively) while the LipidTox signal (green channel) remained deactivated. Following cell tracing, the green channel was activated, and lipid droplet tracing was restricted to distinguishable, individual lipid droplets.

#### Deep learning models

To automate the quantification of large vacuoles, deep learning models were trained using Keras (Chollet 2015) and Tensorflow v2.5 (Abadi et al. 2015) on a NVIDIA RTX 3090 graphics processing unit. Two models were developed. First, a RetinaNet-based model (Lin et al. 2018) was trained to localize CPECs within an image, and then a ResNet-based model was trained to classify CPECs with and without large vacuoles. The first model adopted the RetinaNet architecture as described in Lin et al. (2018) with modifications (see Fig. S3). The model input image size was 256 x 256 pixels, and the number of classes (*K*) and different anchor aspect ratios (*A*) were both reduced to 1. Because all images fed into the model had the same magnification and resolution, only the P3 feature map was passed to the regression and classification subnets. For the second model, hyperparameter tuning on ResNet-based models using keras-tuner (O’Malley et al. 2019) determined that ResNeXt-50 architecture (Xie et al. 2017) with a conv3_depth of 8, conv4_depth of 36, average pooling, and RMSprop optimizer for training had the best result. This ResNeXt model had an input size of 128 x 128 and a fully connected layer output of 2 (CPECs with and without large vacuoles).

Training, validation, and test datasets were developed using the annotations from the manual analysis of vacuoles. For CPEC detection by the RetinaNet model, this included 7502, 1607, and 1607 images with annotated CPECs for training, validation, and test datasets, respectively. For large vacuole classification, this included 118706, 29676, and 37095 CPEC images annotated for the presence of large vacuoles for training, validation, and test datasets, respectively.

Training of the two models was as follows: The RetinaNet model was trained using the Adam optimizer (Kingman & Ba, 2017) with focal loss and Huber loss functions as provided in the jarvis-md python package (https://pypi.org/project/jarvis-md/). Before being passed into the model, images were first standardized to have a pixel mean of 0 and standard deviation of 1. The model was trained with mini-batches of 16 images with on-the-fly data augmentation including random contrast adjustment. During model inference, box predictions were pruned using a non-max suppression threshold of 0.3. The ResNeXt large vacuole classifier model was trained using the RMSprop optimizer (https://www.cs.toronto.edu/~tijmen/csc321/slides/lecture_slides_lec6.pdf) with a categorical cross-entropy loss function. Similar to our RetinaNet training, images were standardized to have a pixel mean of 0 and standard deviation of 1 prior to being passed into the model. The model was trained with mini-batches of 64 images with on-the-fly data augmentation including random flipping, rotation, and contrast adjustment.

All relevant code regarding model architecture setup and training can be found in our Github repository [https://github.com/monukilab/Lipid-storage-paper], and image datasets are available upon request.

## Statistics

Statistical analysis of postmortem data was done under the guidance of the UCI Center for Statistical Consulting. Size distribution analysis used GraphPad Prism9 software. SPSS version 28 was used for other statistical tests on postmortem data, which involved mixed linear models with robust errors to account for heteroskedasticity in the data. Statistical analysis of lipid uptake used GraphPad Prism9 software.

## SUPPLEMENTAL TABLES AND FIGURES

**Table S2.**
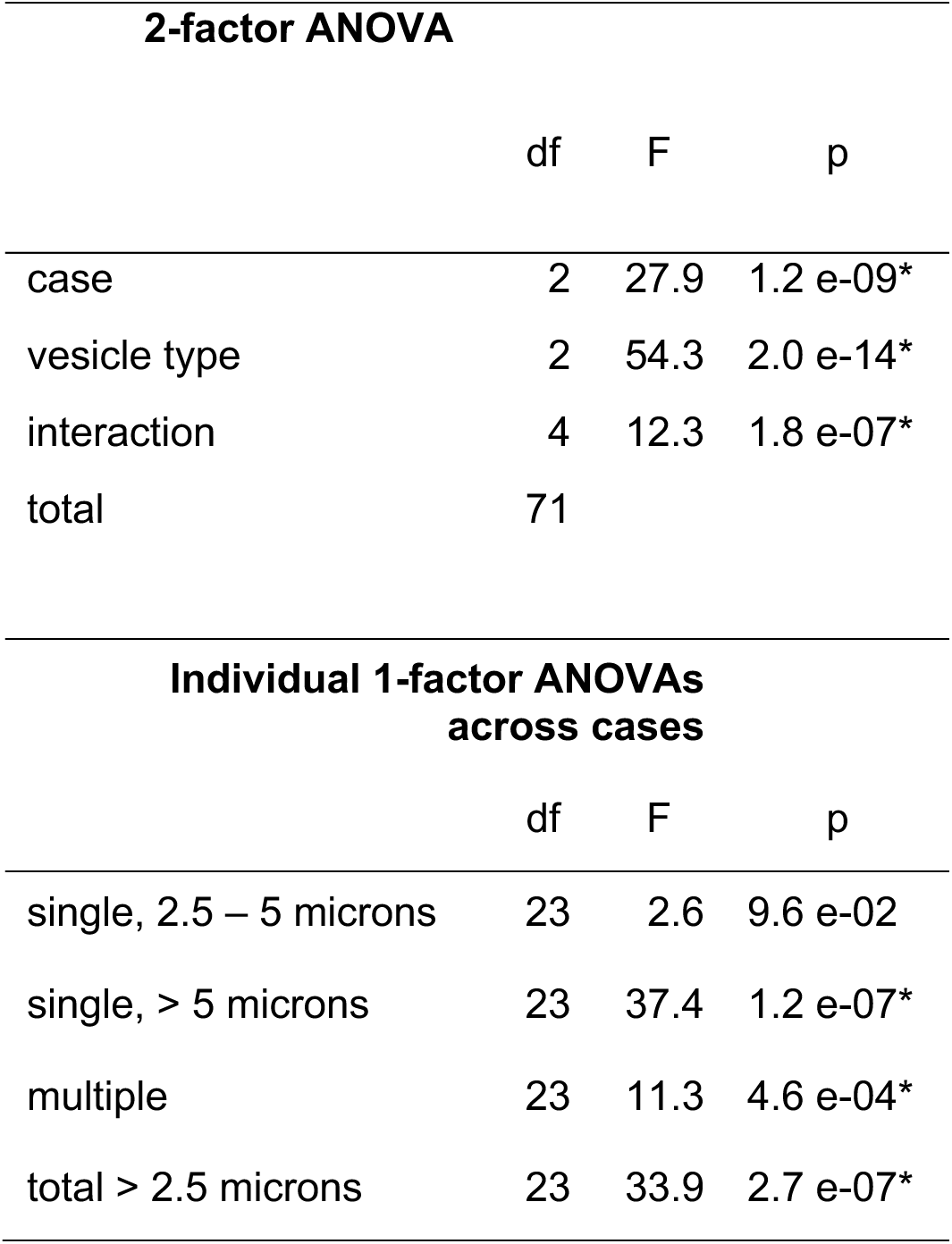
Statistically significant differences (p<0.05, *) across vesicle types and cases despite variance among annotators.

| 2-factor ANOVA |  |  |  |
| --- | --- | --- | --- |
|  | df | F | p |
| case | 2 | 27.9 | 1.2 e-09* |
| vesicle type | 2 | 54.3 | 2.0 e-14* |
| interaction | 4 | 12.3 | 1.8 e-07* |
| total | 71 |  |  |

| Individual 1-factor ANOVAs<br>across cases |  |  |  |
| --- | --- | --- | --- |
|  | df | F | p |
| single, 2.5 – 5 microns | 23 | 2.6 | 9.6 e-02 |
| single, > 5 microns | 23 | 37.4 | 1.2 e-07* |
| multiple | 23 | 11.3 | 4.6 e-04* |
| total > 2.5 microns | 23 | 33.9 | 2.7 e-07* |

**Table S3.** Statistical analysis of size distribution data for each of three annotators (BJ, RS, and NM) showing similar findings of differences and similarities across four different cases. Asterisks denote p < 0.05. Graphical depiction of the raw data can be found in Supplemental Figure 3.

| case | n |  |  | Kruskal-Wallis |  |  |
| --- | --- | --- | --- | --- | --- | --- |
|  | BJ | RS | NM |  | stat | p |
| 783896 | 524 | 499 | 500 | BJ | 211.5 | <0.0001* |
| 763917 | 512 | 523 | 528 | RS | 287.8 | <0.0001* |
| 447764 | 518 | 537 | 509 | NM | 264.3 | <0.0001* |
| 164179 | 510 | 520 | 504 |  |  |  |

| Mann-Whitney U |  |  |  |  |  |  |  |
| --- | --- | --- | --- | --- | --- | --- | --- |
|  | 763917 |  | 447764 |  | 164179 |  |  |
|  | U | p | U | p | U | p |  |
| 783896 | 72506 | <0.0001* | 120926 | 0.0023* | 121588 | 0.012* | BJ |
|  | 56158 | <0.0001* | 94162 | <0.0001* | 95254 | <0.0001* | RS |
|  | 60849 | <0.0001* | 99498 | <0.0001* | 102268 | <0.0001* | NM |
| 763917 |  |  | 79688 | <0.0001* | 78571 | <0.0001* | BJ |
|  |  |  | 87398 | <0.0001* | 88066 | <0.0001* | RS |
|  |  |  | 78099 | <0.0001* | 81172 | <0.0001* | NM |
| 447764 |  |  |  |  | 128328 | 0.430 | BJ |
|  |  |  |  |  | 136757 | 0.564 | RS |
|  |  |  |  |  | 124351 | 0.400 | NM |

| Kolmogorov-Smirnov |  |  |  |  |  |  |  |
| --- | --- | --- | --- | --- | --- | --- | --- |
|  | 763917 |  | 447764 |  | 164179 |  |  |
|  | D | p | D | p | D | p |  |
| 783896 | 0.354 | <0.0001* | 0.152 | <0.0001* | 0.110 | 0.0038* | BJ |
|  | 0.466 | <0.0001* | 0.245 | <0.0001* | 0.199 | <0.0001* | RS |
|  | 0.442 | <0.0001* | 0.228 | <0.0001* | 0.157 | <0.0001* | NM |
| 763917 |  |  | 0.350 | <0.0001* | 0.338 | <0.0001* | BJ |
|  |  |  | 0.311 | <0.0001* | 0.320 | <0.0001* | RS |
|  |  |  | 0.361 | <0.0001* | 0.348 | <0.0001* | NM |
| 447764 |  |  |  |  | 0.081 | 0.071 | BJ |
|  |  |  |  |  | 0.075 | 0.105 | RS |
|  |  |  |  |  | 0.090 | 0.033* | NM |

**Table S4.** Sensitivity, precision, and false discovery rates (FDR) of CPEC detection from three different whole slide images, comparisons between the results of the neural network model with manual data from three different annotators.

| Case | Annotator | Sensitivity | Precision | FDR |
| --- | --- | --- | --- | --- |
| 363485 | RS | 0.848 | 0.888 | 0.112 |
| 185324 | BJ | 0.811 | 0.920 | 0.080 |
| 335909 | SP | 0.867 | 0.805 | 0.195 |

---

**Fig. S1.**
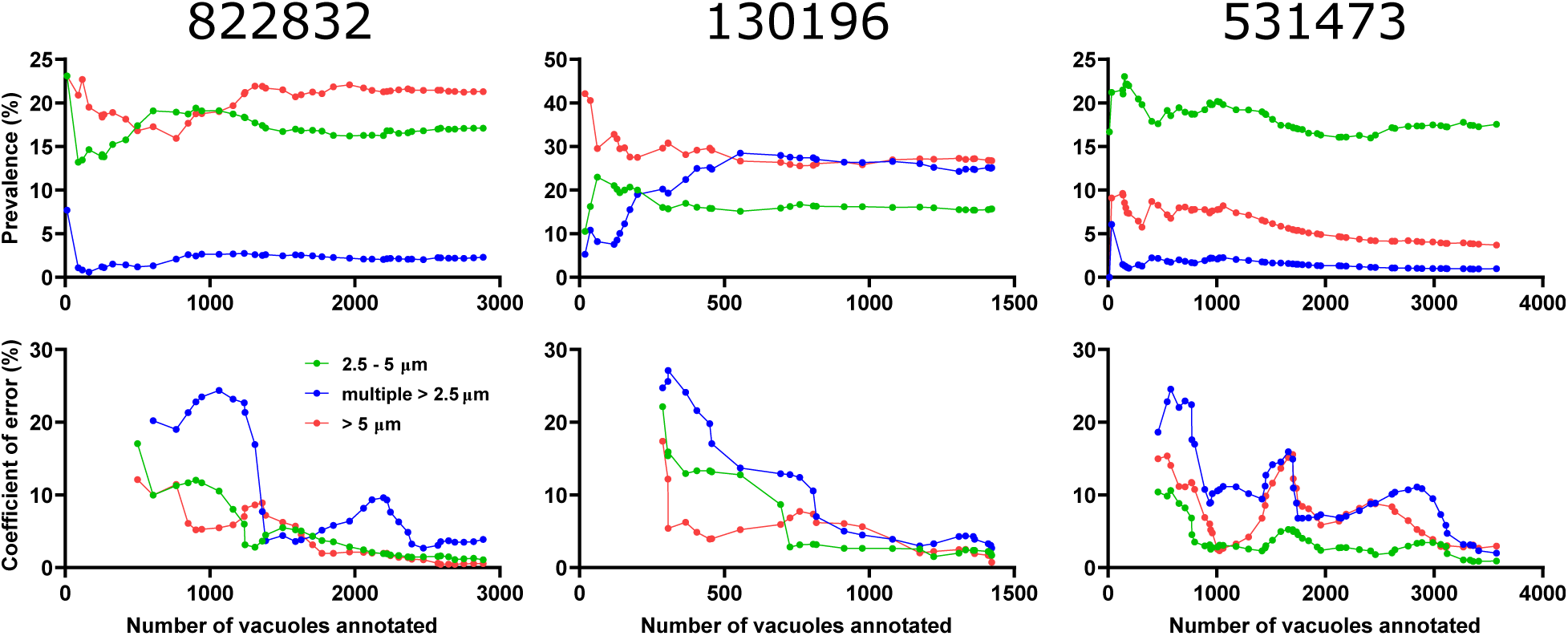
Plots illustrating the method to judge adequate sampling of whole slide images of H&E-stained ChP sections. The whole slide images were divided into equally sized tiles, which were then presented to an annotator in a randomized order. All vacuoles in a tile were assessed, and the estimate of the cumulative prevalence of each vacuole category was calculated at the end of the tile. The top panels show these estimates as a function of the cumulative number of vacuoles annotated at the end of each tile. Starting with the tenth tile, the standard deviation of the estimates across the last ten tiles was expressed as a percentage of the current estimate and plotted as coefficients of error in the bottom panels. Once the coefficients remained below 5% for all categories across ten consecutive tiles (or if any category resulted in a prevalence below 2% when other categories satisfied the criterion), the case was judged as having been adequately sampled, and the values were exported. The three cases in this figure were chosen to illustrate differences in the number of vacuoles required to reach completion, as well as differences in the final prevalence of vacuoles in different categories.

**Fig. S2.**
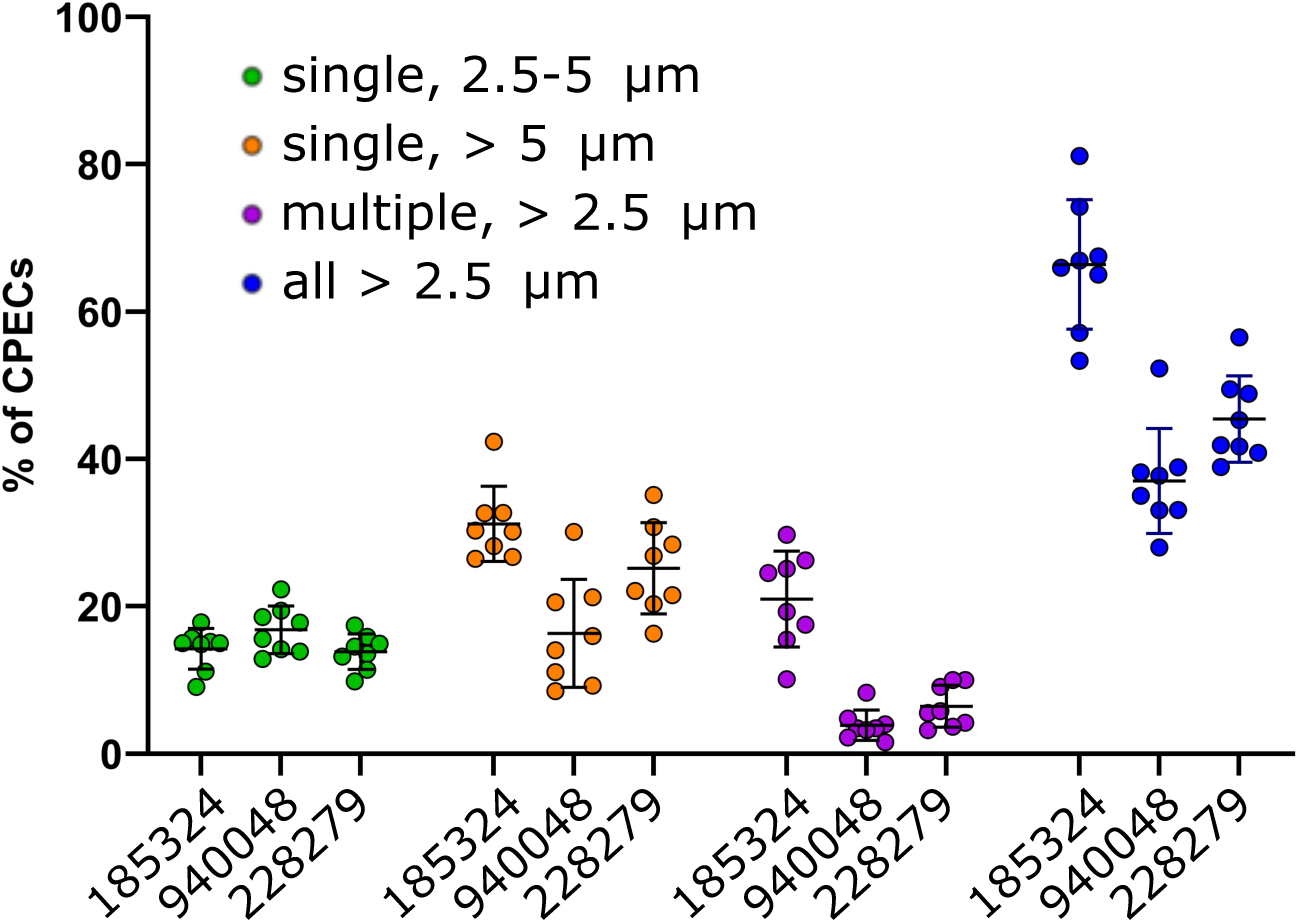
Annotator variance. For three cases, eight different annotators assessed the same tiles to judge observer error in the prevalence of different categories. Error bars denote means and standard deviations across the eight observers. Cases were found to differ significantly (2-factor ANOVA, Suppl. Table 2) despite considerable variance across annotators.

**Fig. S3.**
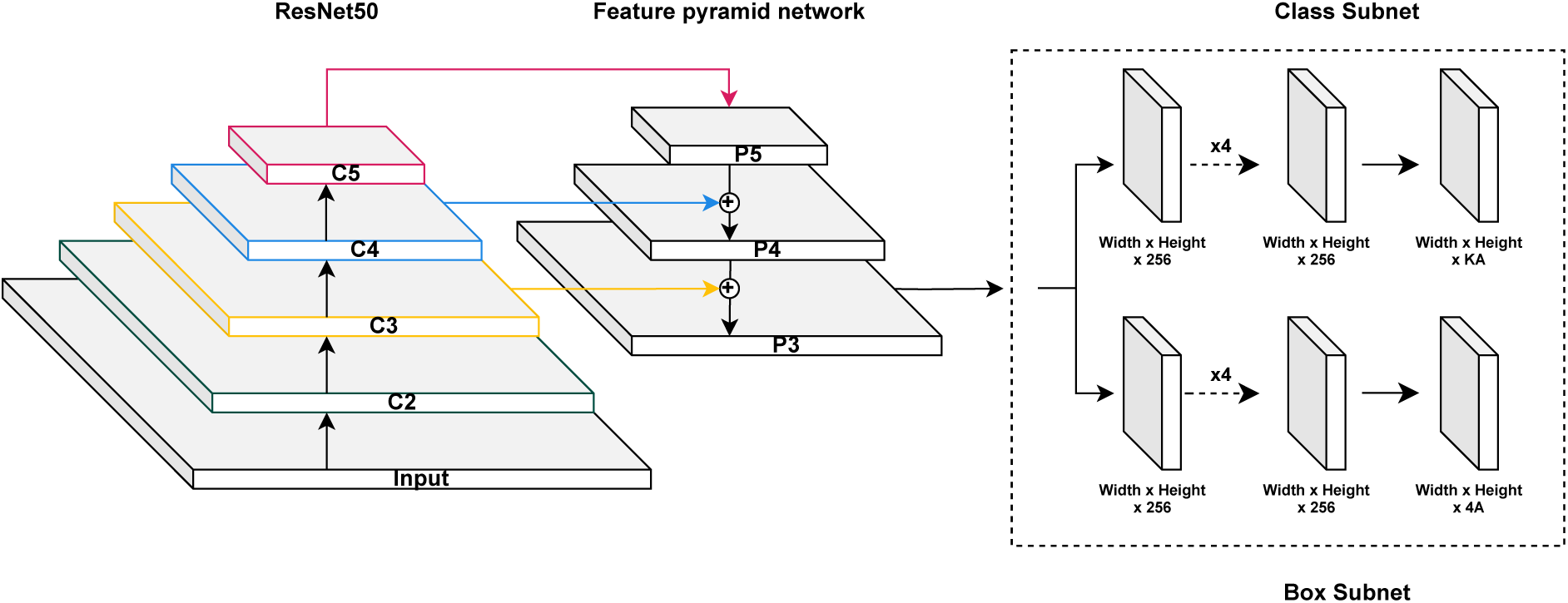
CPEC detection model Architecture. Model architecture diagram showing the modified RetinaNet model used for CPEC detection. The main difference is the use of only the P3 feature map for the class and box subnets.

**Fig S4.**
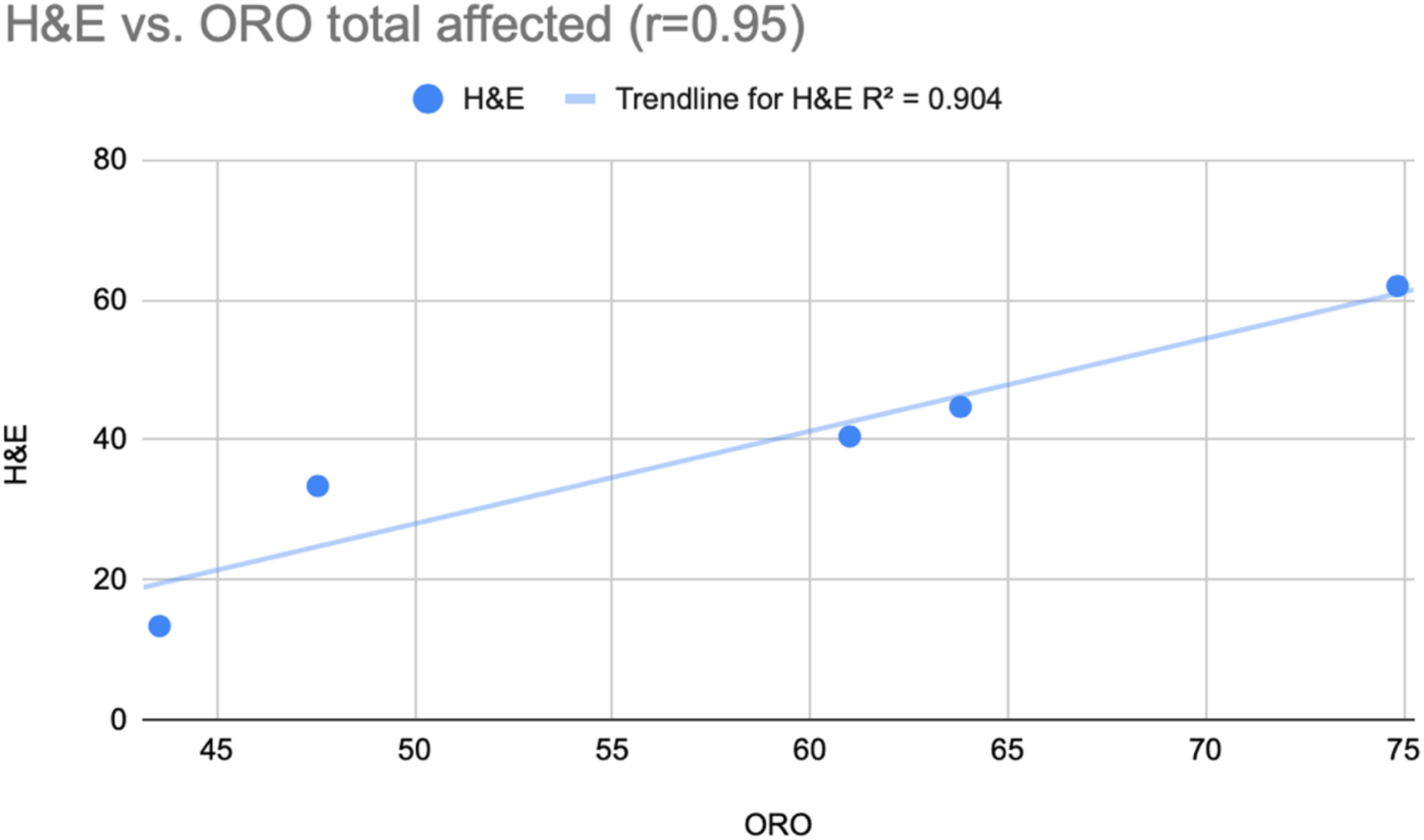
Quantification of percent CPECs affected for five cases using Oil red O-stained sections or separate H&E sections.

**Fig. S5.**
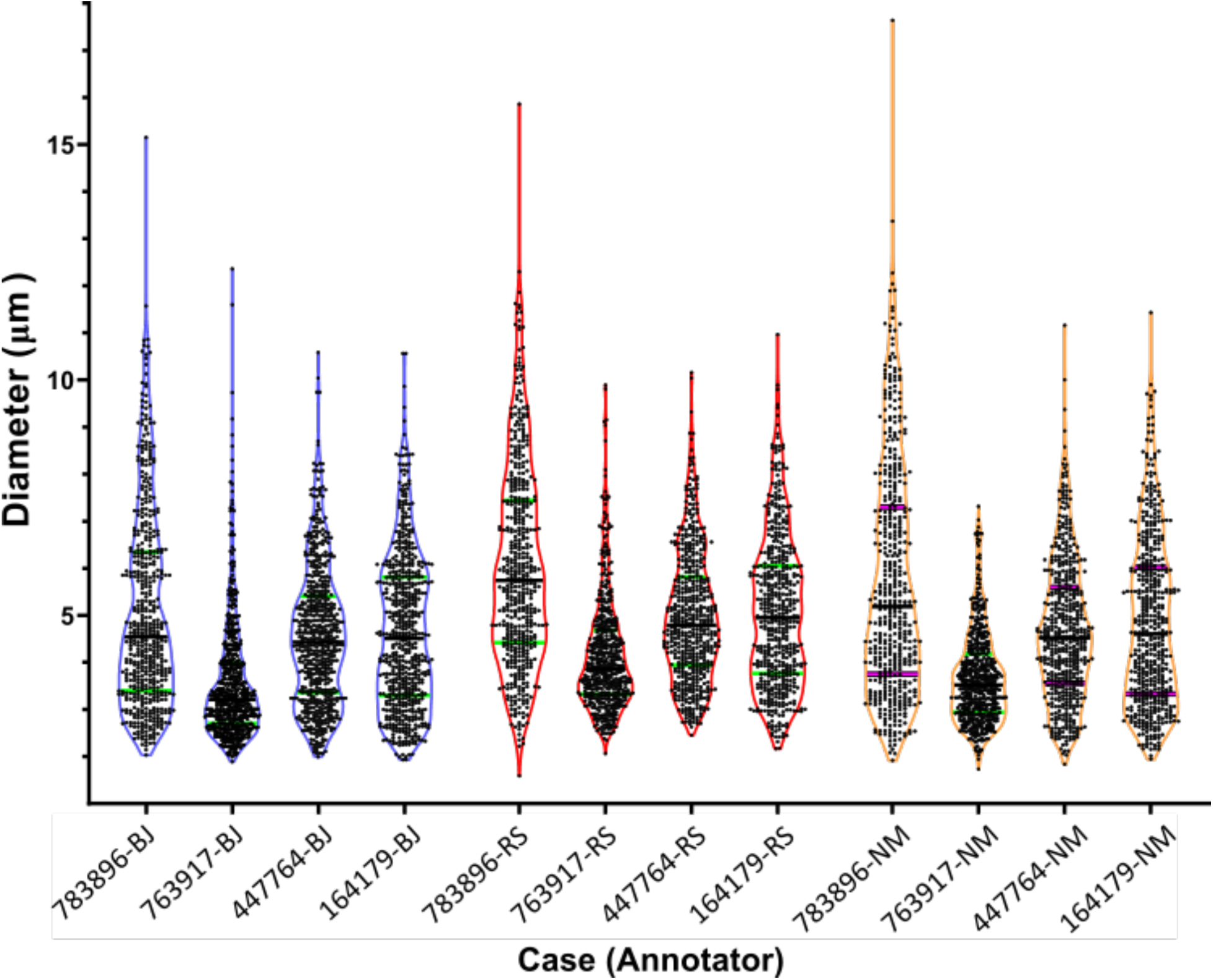
Violin plots showing H&E vacuole size distributions for four cases analyzed independently by three different annotators. Areas of about 500 vacuoles greater than 2.5 µm across their longest axis were measured for each case, and the diameters of circles that would result in those areas were calculated and plotted. Each point represents a vacuole, median sizes are indicated with a black line and quartiles are shown in green. The cases showed similar differences in size distribution across the different annotators, resulting in vacuoles exceeding 10 µm in diameter. Statistical comparisons across cases for each annotator are shown in Supplementary Table 3. Annotators differed mostly in their assessment of vacuoles with the smallest diameters.

**Fig. S6.**
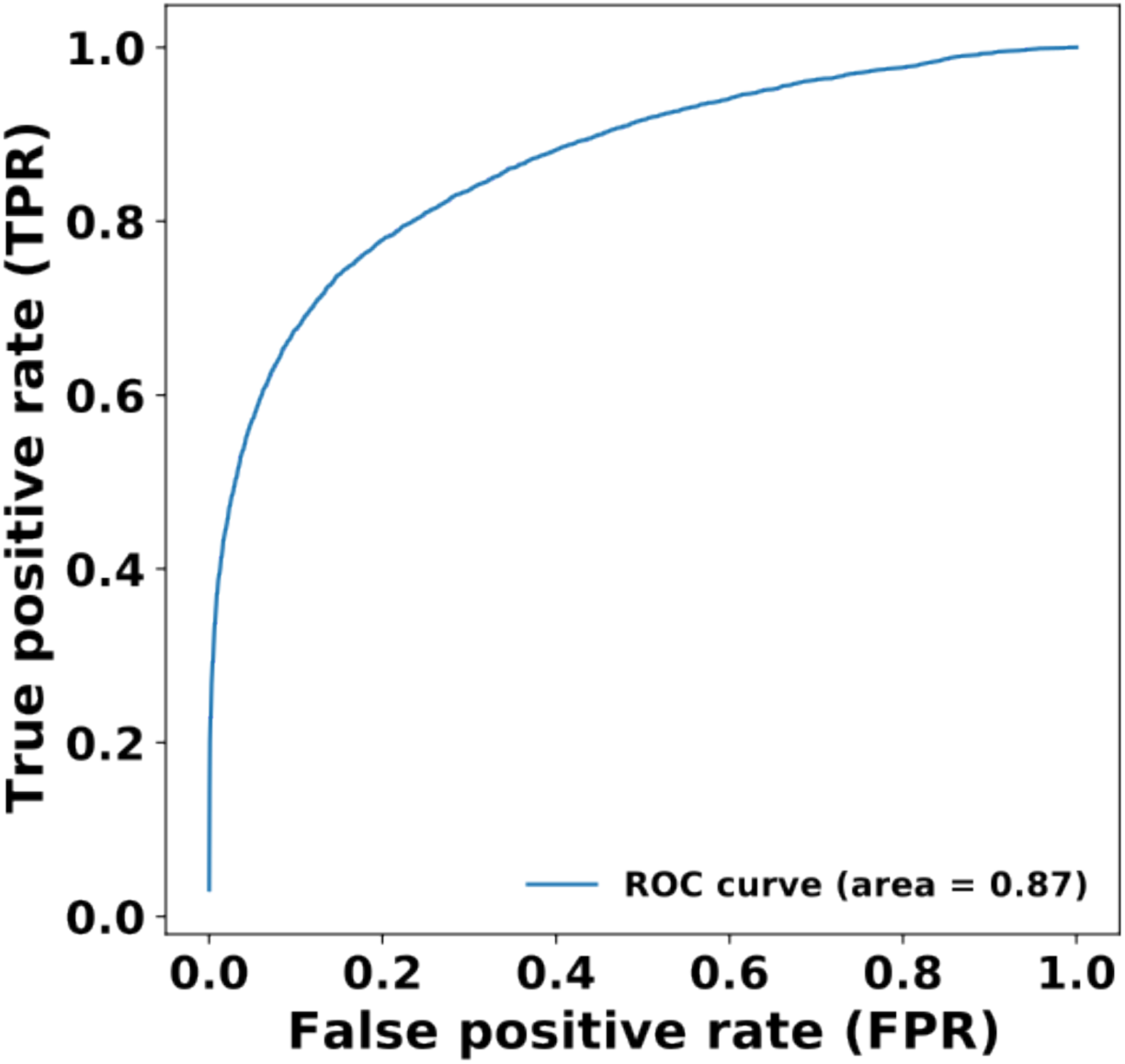
Receiver operating characteristic curve showing performance of the large lipid vacuole classification.

**Fig. S7.**
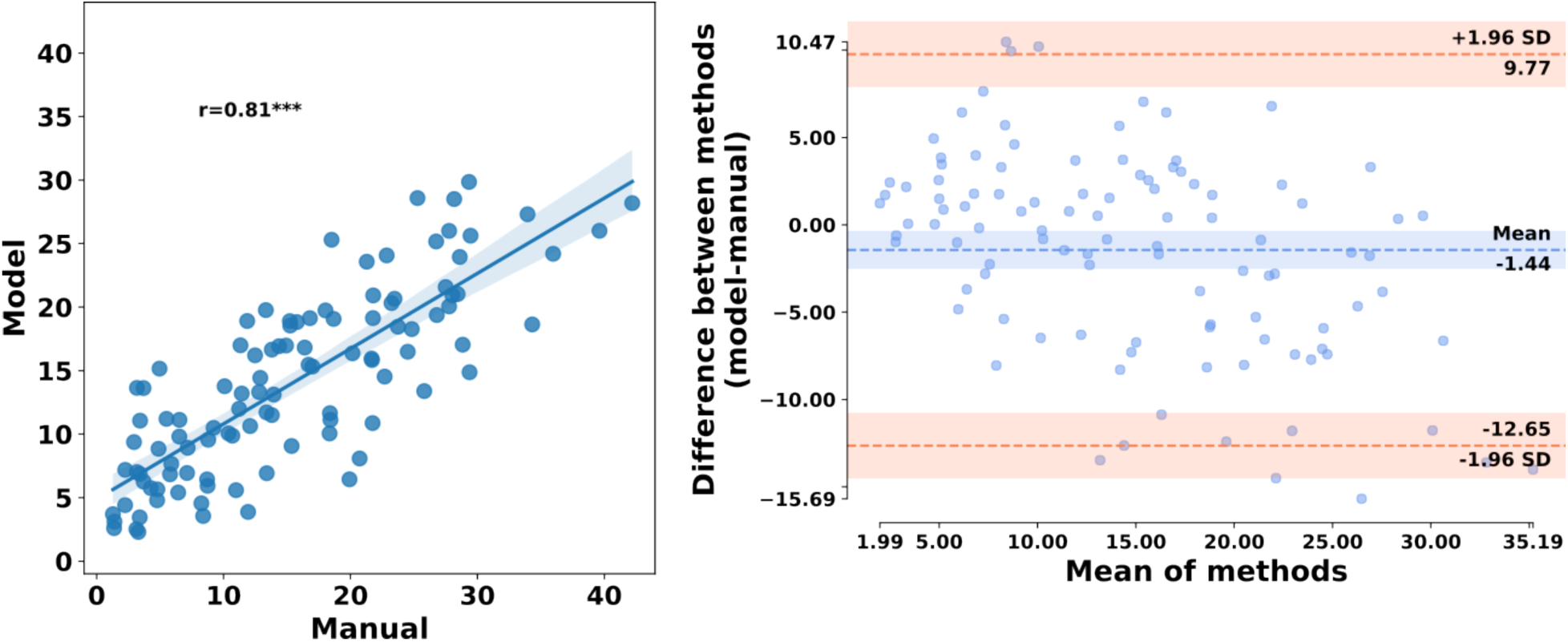
Agreement between manual results and results from the deep learning models. **A)** Correlation analysis comparing model-derived large lipid vesicle data based on entire whole slide images and manual-derived data based on random tile sampling. **B)** Bland-Altman plot comparing model- and manual-derived large lipid vesicle prevalence. In general, the model slightly underestimated manual data (blue shaded line), with a slight proportional bias toward greater underestimation at higher percentages of cells affected, but most individual values fell within the 95% confidence interval of agreement (area between peach shaded lines).

**Fig. S8.**
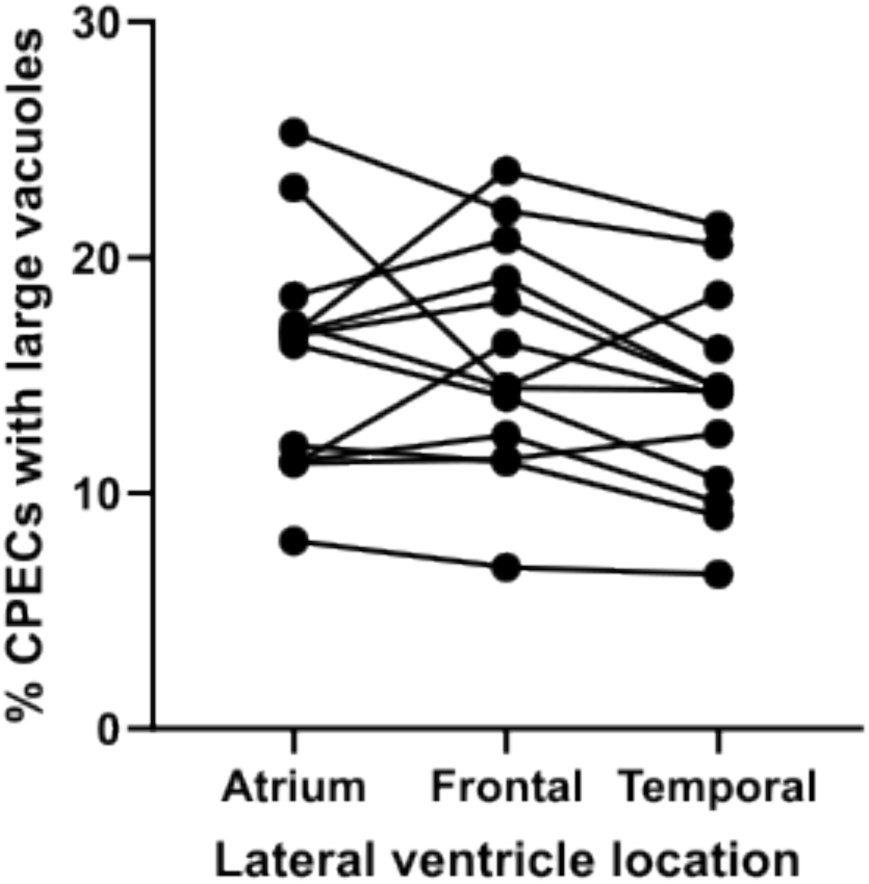
No systematic differences in large lipid vacuoles across locations within the lateral ventricle. A total of 13 cases for which samples were available from the frontal and temporal horns of the lateral ventricle as well as from the atrium were subjected to analysis by our convolutional neural network approach. Individual plot symbols represent analyses of one whole slide image. Sections involving different ventricle locations from the same case are connected by line segments. Whereas there were significant differences among the cases, there were no significant differences across locations (repeated measures ANOVA, F2=2.592, p=0.096).

**Fig. S9.**
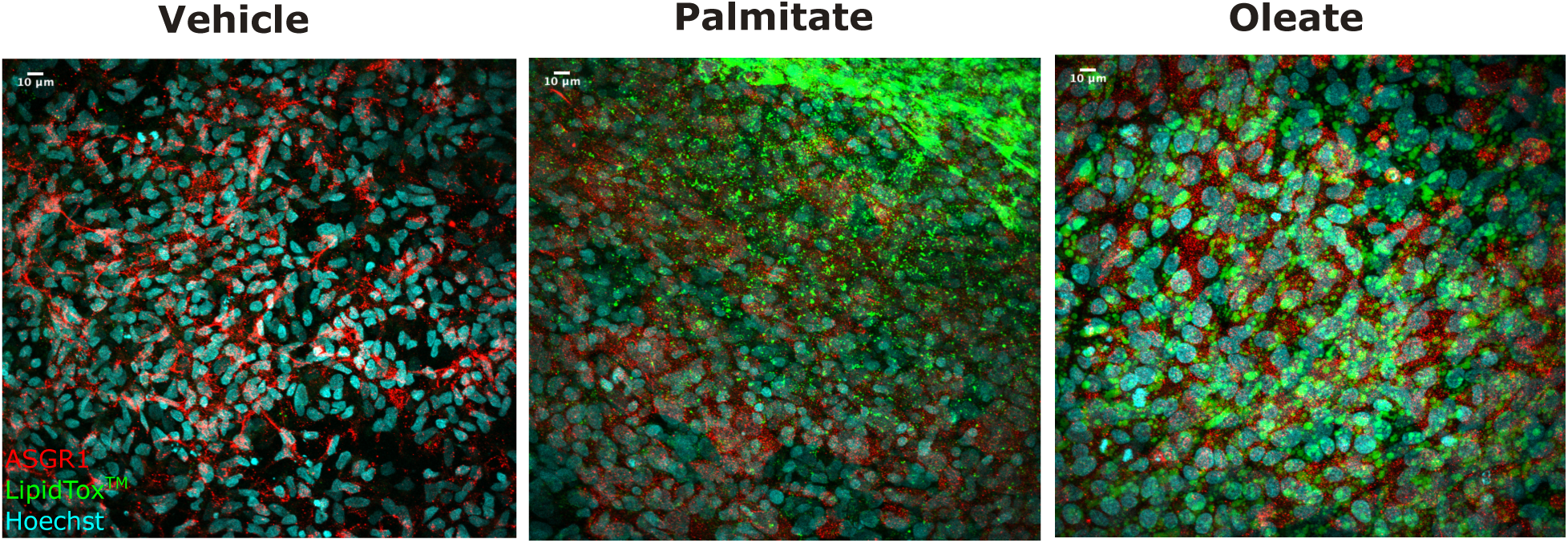
Confocal images of iPSC-derived hepatocytes demonstrating a lipid treatment effect compared to vehicle (BSA only). Scale bar: 10µm. Stained for ASGR1 (red), LipidTOX^TM^ (green), and Hoechst (blue).

## SUPPLEMENTAL FILE

**Supplemental Table 1:** Individual case data

